# Mobile Brain/Body Imaging of cognitive-motor impairment in multiple sclerosis: deriving EEG-based neuro-markers during a dual-task walking study

**DOI:** 10.1101/743328

**Authors:** Pierfilippo De Sanctis, Brenda R. Malcolm, Peter C. Mabie, Ana A. Francisco, Wenzhu B. Mowrey, Sonja Joshi, Sophie Molholm, John J. Foxe

## Abstract

Individuals with a diagnosis of multiple sclerosis (MS) often present with deficits in the cognitive as well as the motor domain. The ability to perform tasks that rely on both domains may therefore be particularly impaired. Yet, behavioral studies designed to measure costs associated with performing two tasks at the same time such as dual-task walking have yielded mixed results. Patients may mobilize additional brain resources to sustain good levels of performance. To test this hypothesis, we acquired event-related potentials (ERP) in thirteen individuals with MS and fifteen healthy control (HC) participants performing a Go/NoGo response inhibition task while sitting (i.e., single task) or walking on a treadmill (i.e., dual-task). In previous work, we showed that the nogo-N2 elicited by the cognitive task was reduced when healthy adults are also asked to walk, and that nogo-N2 reduction was accompanied by sustained dual-task performance. We predicted that some MS patients, similar to their healthy peers, may mobilize N2-indexed brain resources and thereby reduce costs. Somewhat to our surprise, the HC group performed the Go/NoGo task more accurately while walking, thus showing a dual-task benefit, whereas, in line with expectation, the MS group showed a trend towards dual-task costs. The expected nogo-N2 reduction during dual-task walking was found in the HC group, but was not present at the group level in the MS group, suggesting that this group did not modulate the nogo-N2 process in response to higher task load. Regression analysis for the pooled sample revealed a robust link between nogo-N2 reduction and better dual-task performance. We conclude that impaired nogo-N2 adaptation reflects a neurophysiological marker of cognitive-motor dysfunction in MS.

## INTRODUCTION

How do individuals with mobility and cognitive limitations leverage their brain resources to most effectively organize their behavior as they ambulate through a complex and ever-changing environment? This question captures a central issue faced by individuals with neurological diseases such as multiple sclerosis (MS). Limitations may be overcome through recruitment of additional brain regions, at least during early stages of MS, with neuroimaging studies showing that recruitment of new brain regions or more extensive recruitment of typically engaged regions was associated with better cognitive performance in MS^1–4^. How brain resources are effectively deployed during ambulation is less clear due to the requirement of most imaging approaches to remain stationary during recordings. More recent studies acquiring electroencephalographic (EEG) recordings during walking and applying advanced signal processing to correct for motion and muscle artefacts have demonstrated the feasibility of obtaining clean electro-cortical signals during locomotion^5–10^. Here, we employed an EEG-based dual-task walking design to determine whether some individual with MS efficiently deploy cortical resources to maintain a high level of performance during multitasking.

One common approach to investigate cognitive-motor coupling in the laboratory setting is the dual-task walking paradigm ^11–13^. Participants are asked to walk while simultaneously engaging in a cognitive task. Changes in gait, such as decreased walking speed are measured relative to a walking-only task. Similarly, changes in cognition such as increased error rates are measured relative to a cognitive-only task (e.g. participants sit or stand while performing the cognitive task). Relative decrements in performance are called dual-task costs (DTC) and may be suggestive of cognitive-motor interference (CMI). Individuals with MS show robust costs in the form of reduced walking speed and/or poorer cognitive performance as they engage in a secondary task^14–17^. Yet, it is not clear whether MS patients show CMI beyond the costs seen in healthy control participants. A recent meta-analysis of thirteen dual-task walking studies reported small overall effect sizes, and that only 7 out of 13 reported greater costs in individuals with MS relative to individuals without MS^18^. Similarly, studies probing associations between DTC and measures of disability, cognition, and disease duration in MS have produced mixed results^19^. Investigating the neural underpinnings of DTC may provide new insights, possibly showing that individuals with MS with no or equal cost to their healthy counterparts do so by bringing additional brain resources to bear, much as has been seen in fMRI studies, albeit in the absence of ambulation^1–3,20^.

The little that is known about the neural underpinnings of dual-task costs during ambulation in MS^20–23^ is based on functional near infrared spectroscopy (fNIRS) recordings of hemodynamic activity over frontal cortex during walking. Hernandez and colleagues found greater elevation in oxygenation levels over prefrontal regions during Walking While Talking compared to Normal Walking in individuals with MS relative to healthy controls^20^. There was no difference in walking DTC between groups. This is consistent with MS patients compensating for neural degeneration by increasing neural activation to sustain a high level of performance^20^. In contrast, this same group found smaller increases of PFC activation in MS relative to healthy controls^22^ when having participants talk and walk on a 10-cm wide virtual balance beam, which is more demanding and taxes balance. Differences in dual-task complexity may have contributed to differences in their results.

A relatively novel approach known as Mobile Brain/Body Imaging (MoBI) facilitates the integration of electroencephalographic (EEG) recordings and gait kinematics with high millisecond time resolution while participants engage in dual-task walking behaviors^6,24,25^. We previously applied MoBI to measure dual-task effects on gait and the event-related potentials (ERPs) associated with a cognitive task in healthy younger and older adults. Participants performed a Go/NoGo task, which requires overcoming a pre-potent response established by frequent Go stimuli to inhibit response execution to infrequent NoGo stimuli. The ERP response to nogo-trials (requiring response inhibition) compared to go-trials (response execution) is associated with increased frontal N2/P3 amplitudes, with N2 and P3 being negative- and positive-going potentials peaking about 200ms and 300ms after stimulus presentation, respectively^26,27^. Healthy adults showed ERP and gait modulations accompanied by maintained Go/NoGo performance under dual-task load. More specifically, we showed that the nogo-N2 was reduced during walking compared to sitting. In addition, younger adults increased their stride time, thereby making fewer, longer steps, while performing the response inhibition task. In contrast, older adults showed no N2 modulation, no change in stride time, and a decrement in their Go/NoGo performance under dual-task load^28^. We concluded that the nogo-N2 modulation reflected flexible cortical resource allocation under increased task load. Here, we tested the hypothesis that some individuals with MS, similar to their healthy peers, show nogo-N2 modulation linked to better dual-task performance. Establishing neuro-markers of flexible resource allocation during ambulation will provide novel insights into the neural network underlying real-world multitasking difficulties in MS and inform development of targeted interventions aimed at mitigating these issues.

## METHODS

### Participants

Thirteen individuals with relapsing-remitting MS (10 females) were recruited through referrals from a neurologist at Jacobi Medical Center, ads posted on the National MS Society webpage as well as through announcements by leaders of MS support groups organized by the New York Chapter of the National Multiple Sclerosis Society. Individuals, who expressed interest in participating in our study were interviewed over the phone to determine date and type of MS diagnosis. Fifteen healthy controls (9 females) were recruited using flyers, ads, and a laboratory database. Prior to coming in, all volunteers completed a web-based survey to screen for general and mental health. Participants were invited for two visits. The first visit was used to conduct neuropsychological testing and the second visit to perform the EEG study. The first visit lasted approximately 2 hours. The second visit lasted approximately 3.5 hours with .5 hours for capping, 1.5 for data collection, and 1.5 hours for periods of rest to mitigate fatigue. Written consent was required from all participants according to a protocol approved by the institutional review board at Einstein and compliant with the tenets of the Declaration of Helsinki.

#### Demographic and neuropsychological assessment

The following demographic information was obtained from both healthy controls and MS participants: age, gender, and level of education. Participants with MS filled out the Patient-determined Disease Steps (PPDS), which is a self-assessment scale of disease status assessing mobility and daily activities limitation^29^. Cognitive function was evaluated in both groups using the Minimal Assessment of Cognitive Function in MS (MACFIMS), a battery of seven neurological assessments. These include the Paced Auditory Serial Addition Test version 3 (PASAT-3), Symbol Digit Modalities Test (SDMT), California Verbal Learning Test, second edition (CVLT-II), Brief Visuospatial Memory Test – Revised (BVMT-R), Delis-Kaplan Executive Function System (D-KEFS) Sorting Test, Judgment of Line Orientation Test (JLO) and the Controlled Oral Word Association Test (COWAT). These tests assess processing speed/working memory, new learning and recent memory, spatial processing and higher-level executive function, which are commonly used to probe cognitive domains affected in MS^30^. Leg function/ambulation was also measured using the Timed 25-Foot Walk Test and arm/hand function was measured using the 9-Hole Peg Test (9-HPT) to test for mobility differences between healthy controls and MS participants. Finally, we computed the Multiple Sclerosis Functional Composite (MSFC) score based on the Timed 25-Foot Walk, the 9-HPT, and the PASAT-3. The MSFC is a standardize score based on the National Multiple Sclerosis Society’s Clinical Outcomes Assessment Task Force database^31,32^. In two cases, neuropsychological testing was not administered. In total, 14 HCs and 12 MS participants completed the neuropsychological testing battery.

### Stimuli and task

Participants performed a speeded visual Go/No-Go paradigm using images from the International Affective Picture System (IAPS), a database of photographs with normative ratings of emotional status^33^. Only photographs that are classified as affectively neutral or positive were included. Images were presented centrally for 600ms with a random stimulus-onset-asynchrony (SOA) ranging from 800 to 1000ms. Stimuli were presented using Presentation software version 14.4 (Neurobehavioral Systems, Albany, CA, USA) and projected (InFocus XS1 DLP, 1024 x 768 pxl) onto a black wall. On average, images subtended 28° horizontally by 28° vertically. Participants performed the response inhibition task by quickly and accurately clicking a wireless computer mouse button in response to the presentation of each image, while withholding button presses to the second instance of any picture repeated twice in a row. The probability of Go and No-Go trials was 0.85 and 0.15, respectively. Participants completed an average of 12 blocks (each approximately 4 minutes) consisting of three experimental conditions presented in a pseudorandom order: five blocks performing the response inhibition task while sitting, five blocks performing the response inhibition task while walking, and two blocks walking without performing the task. All participants took part in a practice block before undertaking the main experiment. Walking blocks were performed on a treadmill (LifeFitness TR-9000) positioned approximately 1.5 meters from the wall onto which the images were projected. Participants were instructed not to prioritize any single subtask (gait versus cognitive task) but to perform both tasks to the best of their ability. To guard against falls, a custom-designed safety harness was worn while walking (see figure 1 in De Sanctis^11^ for an illustration of the apparatus in use). Participants determined their preferred treadmill walking speed at the beginning of the experimental session, then sustained this speed throughout the duration of the recordings. Average walking speed was 3.53 mph for the healthy control group (range: 2.1 – 4.3) and 3.17 mph (range: 2.4 – 4.0) for the MS patient group.

**Figure 1:**
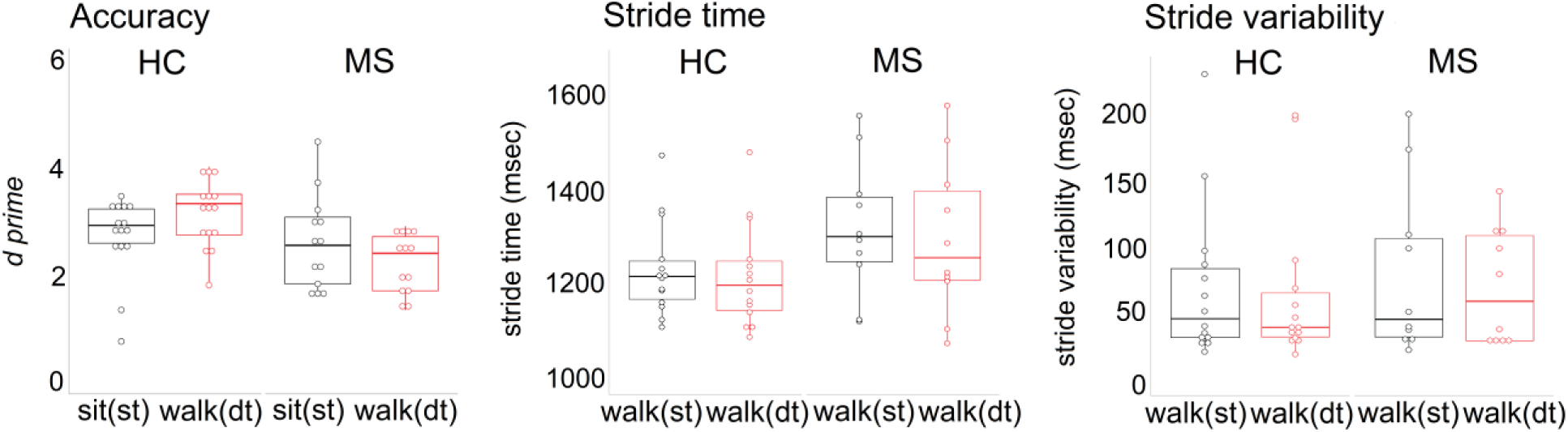
Analysis of Go/NoGo performance (d-prime, left panel) showed a dual-task benefit for HC, while in performance in MS was sustained under dual-task load (st = single task, dt = dual-task). Stride time and stride time variability are displayed for walking-only blocks (black) and walking while performing the cognitive task (red). No group or task load effects were found.

### Gait cycle recording and analysis

Foot force sensors recorded temporal parameters of the gait cycle while participants walked on the treadmill during either uninterrupted walking or while concurrently engaged in the Go/No-Go task. Three sensors (TekscanFlexiForce A201 transducers) were positioned on the sole of each foot: at the center of the back of the heel, the big toe ball and midway along the outer longitudinal arch. These positions enabled the detection of changes in plantar pressure during various stance phases including initial contact, loading response, mid-stance, terminal stance and pre-swing. Force signals were sampled at 512 Hz using an Analog Input Box (BioSemi) connected and integrated via optical fiber with the Biosemi ActiveTwo EEG system. Continuous data were butterworth low-pass filtered at 10Hz, epoched into 10 sec intervals, and normalized against the standard deviation. To assess stride time, we measured peak-to-peak intervals using the force signal derived from a heel sensor (e.g., time of a complete gait cycle is heel contact to next heel contact of that same foot). Automatic peak detection software (MATLAB custom scripts) with one standard deviation as threshold was used to determine if each peak was significantly larger than the data around it. Peak-to-peak intervals were included for further analysis only if the duration to complete a cycle was > 500ms and < 1800ms.

### Event related potential recording and analysis

Scalp recordings were conducted with a 64-channel EEG system (BioSemi ActiveTwo, Amsterdam, The Netherlands), digitized at 512 Hz and bandpass filtered from 0.05 to 100 Hz (24 dB/octave). The BioSemi system uses as reference two electrodes, Common Mode Sense and Driven Right Leg, which form a feedback loop to drive the average potential of the subject (the Common Mode voltage) as close as possible to the ADC reference voltage in the AD-box. Pre-processing and analysis was performed using custom MATLAB scripts (MathWorks, Natick, MA) and EEGLAB^34^. Continuous raw data were re-referenced to CPz, and filtered from 0.5 to 40 Hz to remove low frequency drift and high frequency noise. All blocks across conditions (i.e. sitting or walking) were appended and noisy channels were automatically removed by detecting channels with flat lines (>8 sec.), correlation between neighboring channels < 0.4, and values of line noise exceeding signal by eight standard deviations. In addition to the automated rejection, data were visually inspected and additional channels were excluded if artefacts were present over extended periods of time (~50 sec.). On average, 6 (ranging from 3 to 13) EEG channels were removed due to excessive noise. The remaining channels were re-referenced to a common average reference and visually inspected for prominent artifacts. Next, individual participant data were decomposed using an Independent Component Analysis and components identified as eye movement activity were removed^6,35^. Furthermore, epochs automatically identified as artifactual based on spectrum thresholding were excluded. Finally, excluded channels were re-inserted using a spherical interpolation. We computed epochs time-locked to stimulus presentation with an 800ms post-stimulus period and a 50ms pre-stimulus baseline for Go trials during which the participant successfully responded (Hit trials) and No-Go trials during which the participant successfully withheld a response (Correct Rejection trials [CRs]). Incorrect trials were excluded from the analysis. The average numbers of accepted trials for HC participants were 622 (go) and 52 (nogo) while sitting and 704 (go) and 71 (nogo) during walking. For MS participants, mean of accepted trials were 687 (go) and 57 (nogo) while sitting and 776 (go) and 69 (nogo) during walking.

#### N2amplitude

The N2 ERP component associated with successful response inhibition in a Go/No-Go paradigm has been well characterized in previous studies ^27,36–41^. We quantified the N2 component over central scalp site Cz where we previously found dual-task effects to be of maximal strength^11,42^. We identified the N2 amplitude on a subject-by-subject basis by averaging across trials and subsequently applying automated peak detection within a 180ms time window. The time windows were centered on the nogo-N2 grand means peak amplitude measured in HC at 298ms (during sitting) and 298ms (during walking) and in MS at 314ms (during sitting) and 319ms (during walking). To quantify the N2 amplitude on the single trial-level, we used *area-under-the-curve* measurements between two time points spanning a 60ms period applying the trapezoidal method. Time windows were centered on the N2 peak amplitude derived from single-subject ERP. Statistical analysis was based on N2 *area-under-the-curve* obtained on single-trial level.

#### Statistical analysis

##### Go/NoGo performance

Accuracy in the Go/NoGo task is impacted by task strategy^43^, which may vary between and within a participant (i.e., over the course of the experiment). For example, a participant may decide to adopt a liberal response strategy thereby minimizing misses on go-trials, while accruing higher numbers of false alarms on nogo-trials. Alternatively, a participant may decide to adopt a conservative response strategy thereby minimizing the number of false alarms on nogo-trials, while accruing higher numbers of misses on go-trials. The d-prime calculation based on signal detection theory, is a way to assess task accuracy and account for a participant’s response strategy^43^. Calculation of d-prime is based on the accuracy during go-trials and nogo-trials and reflects the ability to distinguish between target (go) and non-target (no-go) stimuli across trials. We performed a two-way repeated measures analysis of variance (ANOVA) with between-subject factor Group (MS *versus* HC) and within-subject factor Task Load (single-*versus* dual-task load) to analyze task accuracy.

##### Gait performance

Two repeated measures ANOVAs for stride time and stride time variability with between-subject factor Group (MS *versus* HC) and within-subject factor Task Load (single-*versus* dual-task load) were performed.

##### ERP

Mixed-effects models were implemented to analyze the EEG data, using the *lmer* function in the lme4 package^44^ in R (Version 3.1.2, ^45^). The advantages of this approach in modeling EEG data have been previously described^46,47^. Allowing for the modeling of both discrete and continuous variables at multiple levels of variation, mixed-effects models are particularly useful when analyzing complex data. Importantly, compared to traditional ANOVA approaches, mixed-effects models are a) more flexible in dealing with unbalanced and missing data; and b) more flexible regarding statistical dependencies arising from repeated measures (present in EEG experiments given that repeated measurements are taken across trials from the same subjects and that there are spatial correlations between adjacent channels). Area under the curve at Cz was the numeric dependent variable. Group (HC = −0.5, MS = 0.5) was a contrast-coded fixed factor, and task load as well as trial type were numeric fixed factors. Trial was added as random factor, along with by-subject slope adjustments for task load^48^. Models were fit using the maximum likelihood criterion. *P* values were estimated using Satterthwaite approximations.

##### Association between d-prime and ERP modulation

To investigate the relationship between performance in the Go/NoGo task and modulation of the nogo-N2 to increasing task load, we computed the differences in d-prime and nogo-N2 between the dual- and single-task conditions. Positive values for d-prime_diff_ and nogo-N2_diff_ reflect relative better performance and stronger nogo-N2 reduction with increasing task load (i.e., from sitting to walking). In the first model, we performed a linear regression analysis with d-prime_diff_ as the dependent variable and nogo-N2_diff_ as the independent variable. A second model included diagnosis and the interaction diagnosis-by-N2 modulation as predictors to test whether the association differs between groups.

#### Demographic and neuropsychological results

Table 1 displays demographic and clinical information. There were no group differences in age, sex, or years of education. Average time passed since receiving MS diagnosis was 7.34 years (ranging from 1 to 18 years).

**Table 1:** Sample characteristics and neuropsychological test scores for participates in the MS and HC groups. Test scores in the MS group revealed robust deficits in processing speed, verbal fluency, working memory, as well as fine and gross motor impairments.

|  | MS (n=13) | HC (n=15) | p-values |
| --- | --- | --- | --- |
| **DEMOGRAPHIC** |  |  |  |
| Age in years | 34.9 | 34.6 | 0.33 |
| Years of Education | 15.4 | 16.1 | 0.93 |
| Gender (f/m) | 10/3 | 9/6 | 0.33 |
| Years passed since receiving diagnosis | 7.3 | -- |  |
| Patient Determined Disease Steps (PDDS) | 1.58 | -- |  |
| **NEUROPSYCHOLOGICAL TESTS** |  |  |  |
| Visual Spatial Memory (BVMTR-Total Recall) | 24.0 | 26.9 | .264 |
| Visual Spatial Memory (BVMTR – delayed recall ) | 9.0 | 10.7 | .064 |
| Visual Spatial Skills (JLO) | 24.4 | 26.1 | .265 |
| Verbal fluency (COWAT) | 33.8 | 45.7 | **.016\*** |
| Word list total recall (CVLT-II) | 53.4 | 62.7 | **.005\*** |
| Word list long delayed recall (CVLT-II) | 11.5 | 13.9 | **.023\*** |
| Symbol Digit Modalities Test (SDMT) | 45.0 | 57.4 | **.014\*** |
| Fine Motoric (9-Hole Peg Test) | .0481 | .0395 | **.005\*** |
| Paced Auditory Serial Addition Test- v.3 (PASAT-3) | 26.1 | 52.7 | **.000\*** |
| Gait (25-Foot Walk Test) | 7.3 | 4.8 | **.009\*** |
| Multiple Sclerosis Functional Composite (MSFC) | -.54 | .49 | **.000\*** |
| Walking Speed in mph (treadmill) | 1.97 | 2.19 | .192 |

There was a significant group difference for the COWAT (*t*_24_ = 2.6, *p* = .016), the CVLT-II Total Recall (*t*_24_ = 2.8, *p* = .009), the CVLT-II Delayed Recall *(t*_24_ = 2.4, *p* = .023), the SDMT (*t*_24_ = 2.6, *p* = .014), and the PASAT-3 (*t*_24_ = 5.4, *p* < .001). Furthermore, group differences indicating fine and gross motor impairments in our MS group were documented with the 9-HPT (*t*_24_ = 3.1, *p* = .005) and Timed 25 ft. Walk Test (*t*_24_ = −2.8, *p* = .009). Finally, we show group differences for the MSFC composite scores *(t*_24_ = 5.7, *p* < .001). These results indicate that overall the MS group performed more poorly across these three measures (i.e., processing speed, fine motor, and gross motor) in comparison to both the HC group as well as in comparison to the National Multiple Sclerosis Society’s Clinical Outcomes Assessment Task Force database population^31,32^.

#### Behavioral results

Figure 1 (left panel) illustrates results obtained for the d-prime analysis, assessing accuracy in the Go/NoGo task for HC and MS groups performing under single-task (sitting) and dual task (walking) load. For d-prime, a two-way repeated-measures ANOVA with factors of Group and Task Load revealed a significant main effect of Group (F_1, 26_ = 4.90, *p* = .036) due to the HC group performing better overall, and a significant Group X Task Load interaction (F_1, 26_ = 9.42, *p* = .005). Follow-up t-tests indicated that the HC group performed significantly better on the response inhibition task as load increased from sitting to walking (*t*_14_ = 2.23, *p* = .043), while for the MS group, performance declined with increased motor load but this comparison did not reach significance (*t*_12_ = 2.15, *p* = .052).

### Gait cycle results

Figure 1 (middle and right panels) illustrates stride time and stride time variability for single task (black) and dual task (red) walking for HC and MS patients. Foot force sensor recordings from one HC and three MS participants were unusable due to technical difficulties, therefore gait cycle results are reported from 10 MS participants and 14 HCs. Overall, both groups exhibited faster stride times when performing the task (F_1, 22_ = 6.85, *p* = .016). However, after including treadmill walking speed as a covariate there were no longer any differences in average stride time. For stride time variability, there were no significant effects.

### Electrophysiological Results

Figure 2 shows grand mean waveforms with vertical standard error bars at each data point at electrode site Cz for participants in the HC (left column) and MS (right column) groups. Grand mean ERP results are reported from 13 HC and 13 MS participants. Electrophysiological data from two HCs were excluded due to excessive movement-related noise. The ERP modulation as a function of task load during the N2 time period in HC is most clearly illustrated in the bottom row of figure 2, which shows the difference waveforms (CR_ERP_ *minus* Hit_ERP_).

**Figure 2:**
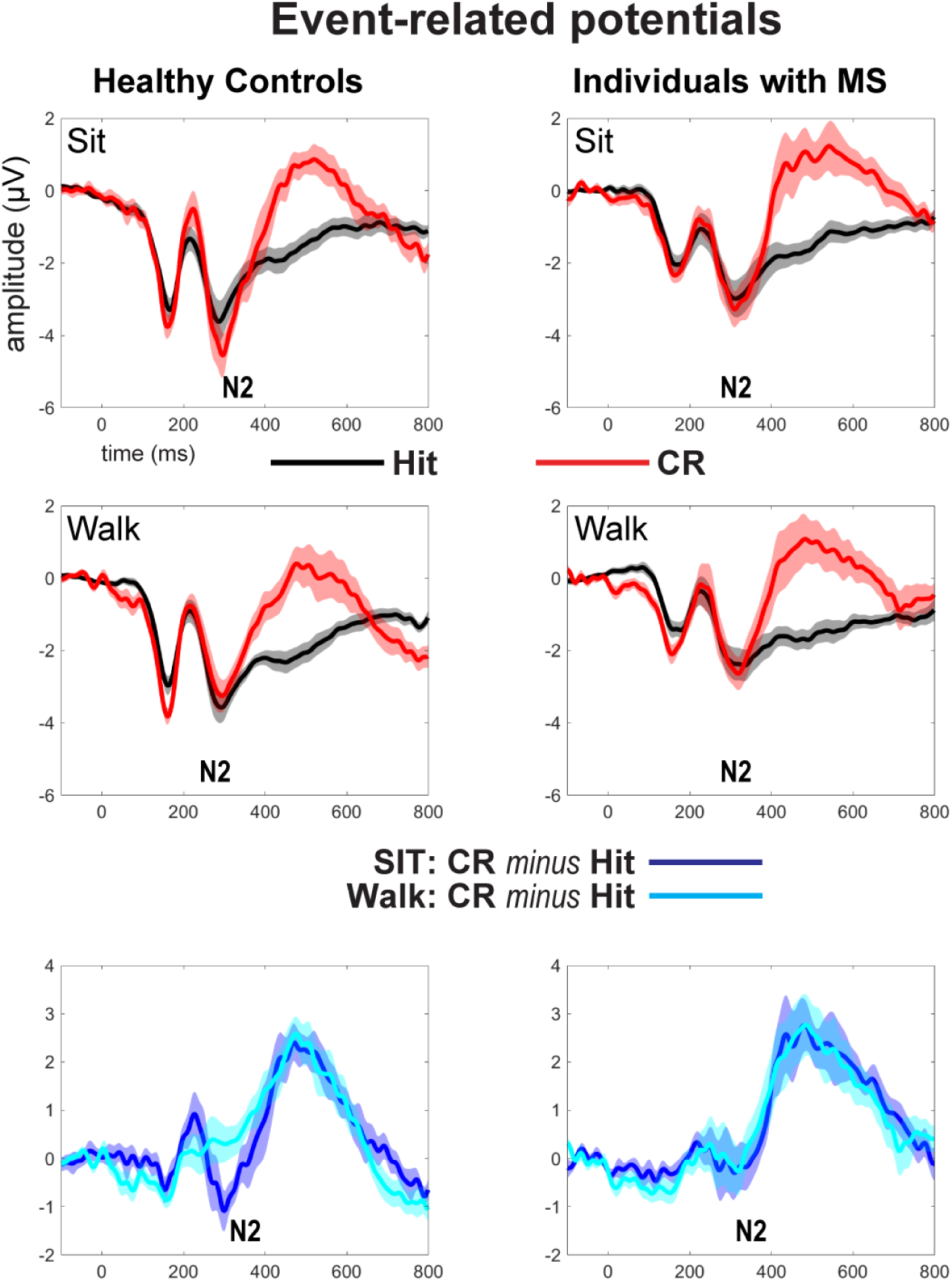
Grand mean ERPs at electrode site Cz for the HC and MS groups performing the Go/NoGo task under single (i.e., sitting; top row) and dual-task (i.e., walking, middle row) load. Difference waves (CRs minus Hits) for sitting and walking conditions are plotted in the bottom row. Difference waveforms most clearly illustrate N2 amplitude modulations as a function of task load in the HC group.

Figure 3 illustrates distribution of single-subject N2 amplitudes as a function of task-load (left column) and trail type (right column). Also, the 2^nd^ and 4^th^ row in figure 3 shows mean differences between conditions and groups (black dots) together with a bootstrap sampling distribution and 95% confidence intervals (vertical back lines).

**Figure 3:**
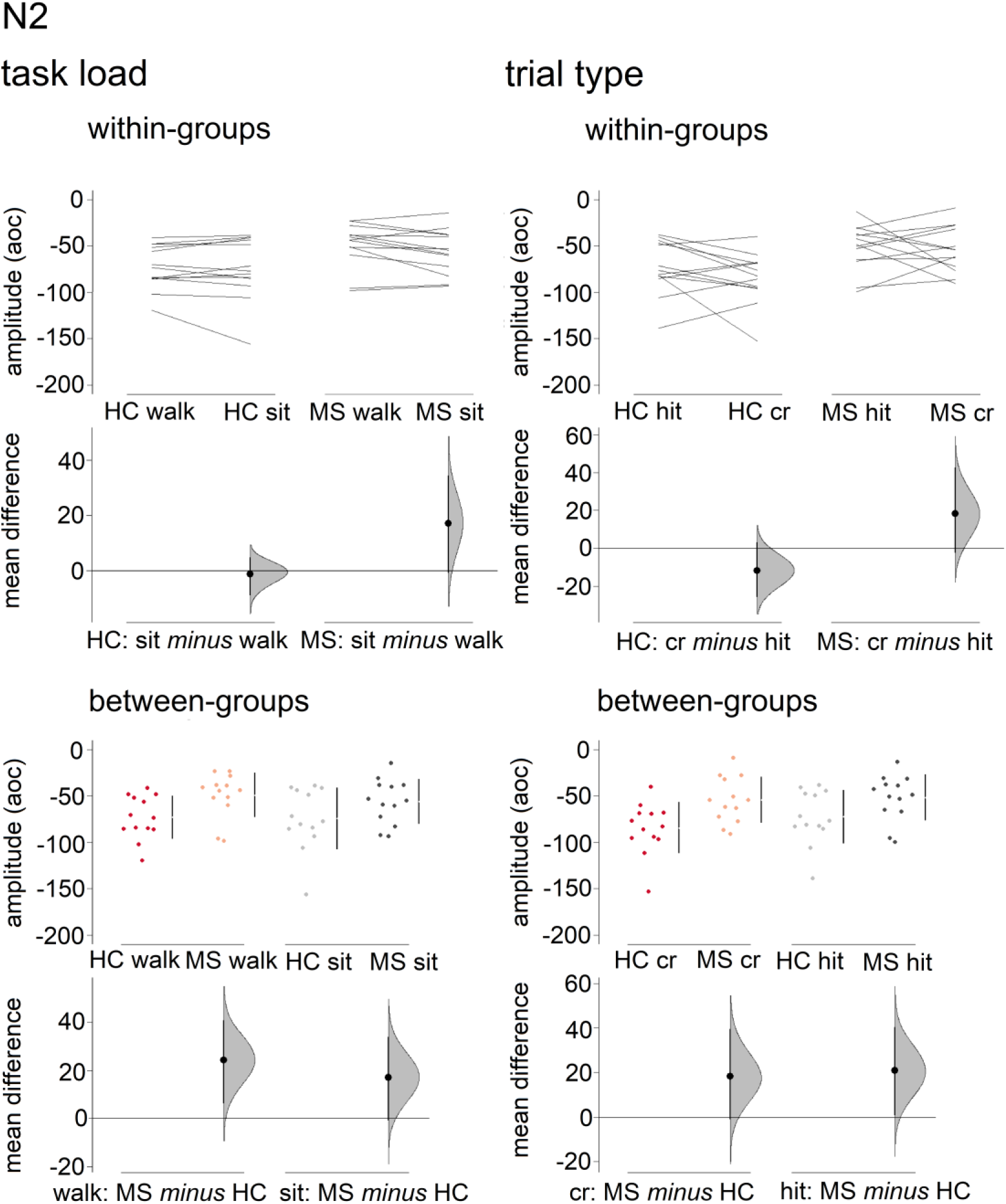
N2 amplitude mean difference for comparisons between groups, task load, and trial types are shown in Cumming estimation plots. The raw data is plotted on the upper axes (summary measurements are displayed as gapped lines to the right of each plot: means are indicated as a gap in the lines, vertical lines represent standard deviation error bars); each mean difference is plotted on the lower axes as a bootstrap sampling distribution. Mean differences are depicted as dots; 95% confidence intervals are indicated by the ends of the vertical error bars ^49^.

Statistical analysis of the N2 revealed the following effects: there was a significant effect of trial type, with correct rejections eliciting a stronger N2 response than hits (*β* = −25.04, SE = 5.26, *p* < .001). The interaction between task load and trial type was also significant (*β* = 24.65, SE = 7.00, *p* < .001): the difference between correct rejections and hits was reduced during dual-task (i.e., walking) compared to single-task load (i.e., sitting). Though no main effect of group was found, the interaction between group and trial type was significant (*β* = 23.91, SE = 7.21, *p* < .001): When compared to the healthy controls, who presented a significantly stronger N2 response for correct rejections than for hits, individuals diagnosed with MS showed less N2 differentiation between correct rejections and hits. Additionally, a three-way interaction between group, task load, and trial type was found to be significant (*β* = −27.96, SE = 9.58, *p* < .01): for the HC, the N2 differentiation between correct rejections and hits was reduced during dual-task compared to single-task load; for the individuals diagnosed with MS the N2 differentiation between correct rejections and hits during single and dual-task load were not statistically different. These interactions can be better appreciated in Figure 4.

**Figure 4:**
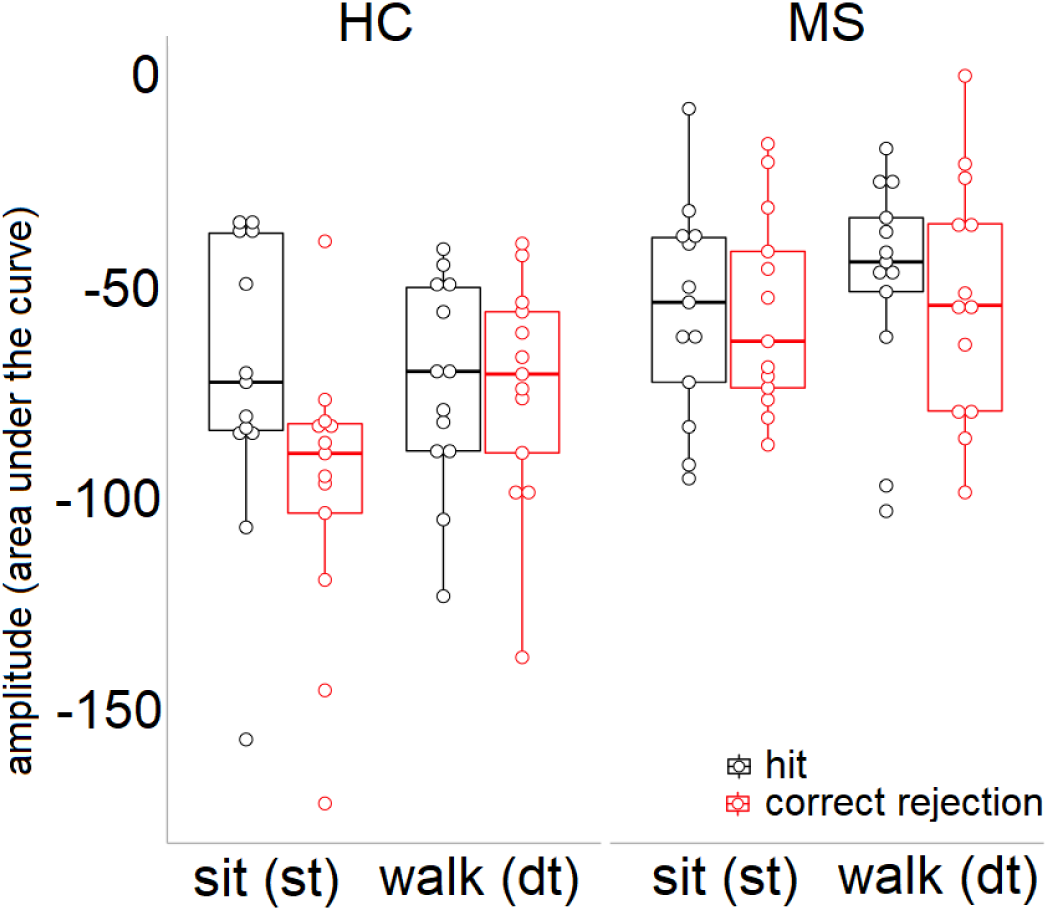
Box plots illustrating N2 amplitude by group and task load. Participants in the HC group showed the N2 modulation as a function of task load.

Figure 5 illustrates the relationship between nogo-N2 modulation and d-prime as a function of task load. Values are plotted as difference measures. More positive values on the x/y axes represent stronger N2/higher accuracy during walking compared to sitting. By pooling participants across both groups, we find a significant correlation (r=−0.44, p=0.02) between stronger nogo-N2 reduction and better Go/NoGo performance during dual-tasking. Yet, after including diagnosis and the interaction, diagnosis-by-N2 modulation, in our model where d-prime is the dependent variable, the interaction term is not significant (p=0.32) and the association between N2 modulation and d-prime was not significant for either diagnosis group (p-values: 0.11 for HC and 0.86 for MS), likely due to insufficient power.

**Figure 5:**
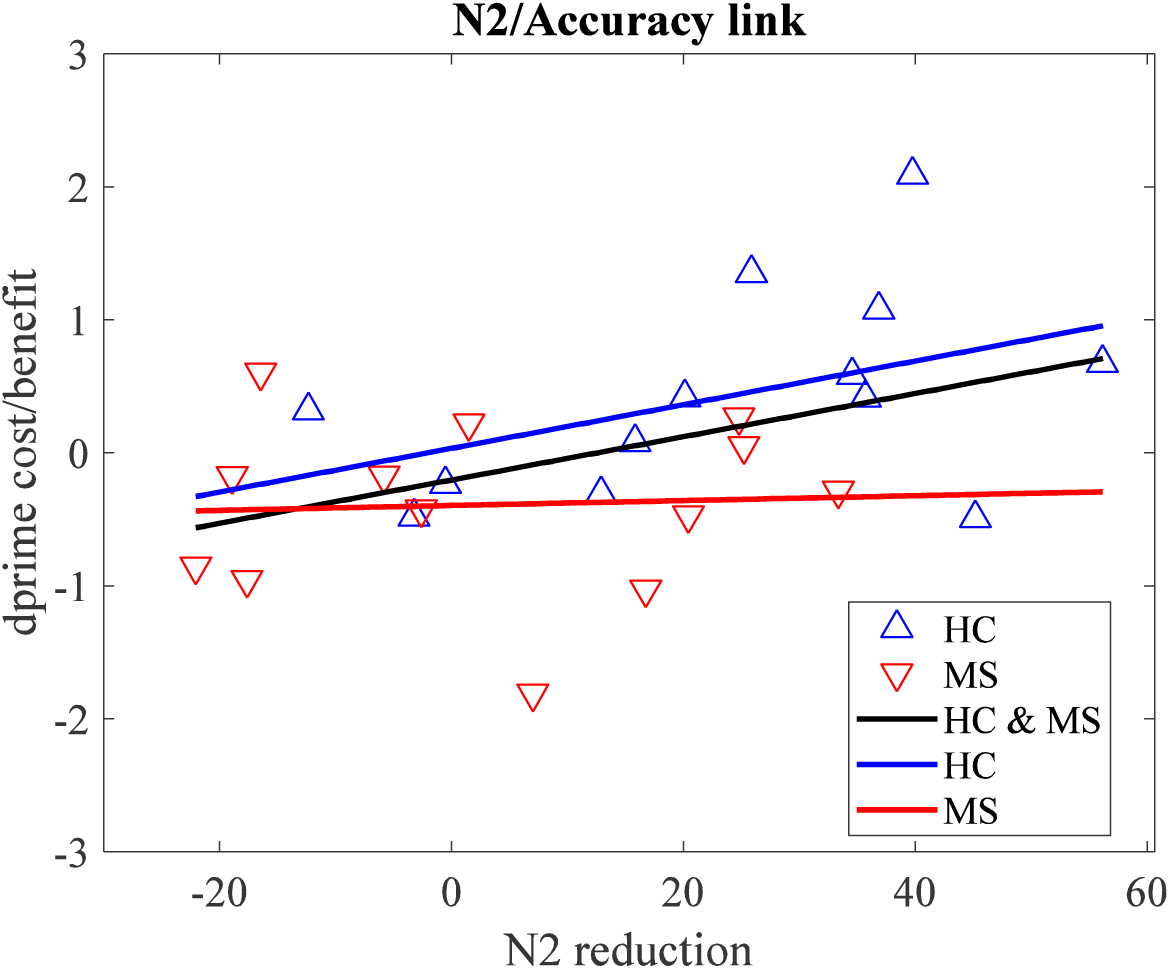
Scatter plot illustrating d-prime and N2 modulation for each participant. Both measures are plotted as difference values between single and dual-task load. As predicted, stronger N2 reduction was association with better Go/NoGo performance during dual-task walking. However, association was not significant after including group and group-by-load interaction as predictor in the model.

## DISCUSSION

We applied a dual-task MoBI approach^5,11,28^ to record brain activity in conjunction with gait and Go/NoGo performance measurements, to gain insights into the cortical correlates of dual-task processing in multiple sclerosis. Based on our prior work, we predicted that some individuals with MS, similar to their healthy control counterparts, are able to adapt information processing to increased dual-task load and thereby reduce costs typically associated with performing two tasks simultaneously ^50,51^. We have previously shown that adapted processing, indexed by load-related N2 modulation, was accompanied by better dual-task performance^28^. We predicted a linear relationship with, stronger N2 modulation being linked to less DTCs. Our goal was to move towards developing neuro-markers of flexible resource allocation during ambulation, which may help design more targeted intervention to mitigate real-world multitasking difficulties in MS.

The Go/NoGo accuracy results are in line with the robust literature on cognitive impairments^52–54^ and also reaffirm the ambiguity of DTC reports in MS^17^. That is, individuals with MS performed worse in the Go/NoGo task, but deficits were not worsened under dual-task load. With regard to our ERP findings, we found group differences with MS patients showing less N2 differentiation between CR and hit trials than HC. Our findings are consistent with the existing literature demonstrating associations between aberrant ERP brain activity and cognitive impairments in MS ^55–57^. The finding of group differences in N2 adaptation to increasing task-load is strikingly similar to our prior MoBI work comparing younger and older adults^28^. The three-way interaction between group, trial type, and task-load confirmed that the N2 differentiation (i.e., N2_CR_ > N2_Hit_) found during single-task load was reduced under dual-task load. The adaptation of nogo-N2 processes to increased task load was present in HC, but not in MS.

Our data do not support our prediction that some individuals with MS adapt nogo-N2 processes to higher task load and thereby minimize DTCs. By pooling participants across both groups, we showed a significant linear association between stronger nogo-N2 reduction and better Go/NoGo performance during dual-tasking. A linear model may, on first sight, suggest that there is a continuum along which participants in both groups deploy the <u>same</u> N2-indexed adaptation process. After including diagnosis and the interaction term diagnosis-by-N2 in our model, however, the association between N2 modulation and d-prime was not significant, possibly due to insufficient power. As can be seen in Figure 5, the significant association for the pooled sample was mostly driven by the HC group. That is, the slopes of the linear function fitting the HC and pooled group were very similar. In contrast, the slope fitting the MS data was close to zero which might suggest that there is no relationship between load-related nogo-N2 modulation and Go/NoGo accuracy.

Differences in adapting the cognitive nogo-N2 to increased motor load point to an impairment of processing and integrating multiple tasks in multiple sclerosis. Relatively higher DTCs may be suggestive of a failure to mobilize additional N2-indexed brain resources during dual-task walking in MS. However, further tests of the N2/d-prime association in MS are required to support the aforementioned conclusion. Of note, recent ERP investigations in MS assessing cognition while participants were stationary (i.e., cognitive-only task) reported evidence in support of compensatory neural functions^55–57^. For example, Lopez-Gongora found larger ERP amplitudes in MS relative to HC, even though groups did not differ in their response inhibition performance.

There are limitations to our study that are important to mention. First, the average time of receiving a diagnosis prior to study participation was 7.3 years and ranged from 1 to 18 years. Including patients with advanced MS may have tempered our goal to identify compensatory functioning, an ability thought to be rather limited to patients during early disease stages. Our sample size consisted of 13 patients. Future studies may increase sample size to enhance the sensitivity to detect dual-task effects on gait and ERP measures. Furthermore, individuals with MS showed a relative decline in Go/NoGo performance under dual-task load, yet the decline was not significant. Robust DTC in the MS group would lend stronger support to our assertion that abnormal nogo-N2 modulation reflects cognitive-motor dysfunction in MS.

In summary, we linked differences in dual-task walking performance to a neurophysiological marker and provided relevant insight into aberrant cortical activity associated with processing and integrating multiple tasks in multiple sclerosis. To our knowledge, this is the first EEG-based dual-task walking study in MS. Cognitive-motor paradigms address more precisely patient-reported multitasking deficits and may therefore improve the sensitivity of patient assessments. Importantly, low cost wireless mobile EEG systems of high signal quality are readily available^58^. We believe that EEG-based measurements of cognitive-motor issues may eventually be sensitive enough to be deployed in the clinical setting to track a patient’s disease progression and test therapeutic efficacy of interventions.

## Acknowledgments

The primary sources of funding for this work was provided by a pilot grant from the National Multiple Sclerosis Society (PP3398), an Exploration - Hypothesis Development Award from the Department of Defense (MS160058), and by a Mentored Research Scientist Development Award from the National Institute on Aging (5K01AG049991). Participant recruitment was performed by the Human Clinical Phenotyping Core at Einstein, a facility of the Rose F. Kennedy Intellectual and Developmental Disabilities Research Center (RFK-IDDRC) which is funded by a center grant from the Eunice Kennedy Shriver National Institute of Child Health & Human Development (NICHD U54 HD090260).

We thank our participants for the willingness to volunteer for this study. We also extend out thanks to neuropsychologist, research technician and interns who helped with assessments and EEG data collection: Dr. Pamela Counts (neuropsychologist), Gregory Peters (technician), Katalina Gomez (intern), and Elizabeth Chernyak (intern).

